# Transmission bottleneck size estimation from *de novo* viral genetic variation

**DOI:** 10.1101/2023.08.14.553219

**Authors:** Teresa Shi, Jeremy D. Harris, Michael A. Martin, Katia Koelle

## Abstract

Sequencing of viral infections has become increasingly common over the last decade. Deep sequencing data in particular have proven useful in characterizing the roles that genetic drift and natural selection play in shaping within-host viral populations. They have also been used to estimate transmission bottleneck sizes from identified donor-recipient pairs. These bottleneck sizes quantify the number of viral particles that establish genetic lineages in the recipient host and are important to estimate due to their impact on viral evolution. Current approaches for estimating bottleneck sizes exclusively consider the subset of viral sites that are observed as polymorphic in the donor individual. However, allele frequencies can change dramatically over the course of an individual’s infection, such that sites that are polymorphic in the donor at the time of transmission may not be polymorphic in the donor at the time of sampling and allele frequencies at donor-polymorphic sites may change dramatically over the course of a recipient’s infection. Because of this, transmission bottleneck sizes estimated using allele frequencies observed at a donor’s polymorphic sites may be considerable underestimates of true bottleneck sizes. Here, we present a new statistical approach for instead estimating bottleneck sizes using patterns of viral genetic variation that arose *de novo* within a recipient individual. Specifically, our approach makes use of the number of clonal viral variants observed in a transmission pair, defined as the number of viral sites that are monomorphic in both the donor and the recipient but carry different alleles. We first test our approach on a simulated dataset and then apply it to both influenza A virus sequence data and SARS-CoV-2 sequence data from identified transmission pairs. Our results confirm the existence of extremely tight transmission bottlenecks for these two respiratory viruses, using an approach that does not tend to underestimate transmission bottleneck sizes.

## Introduction

In viral infections, transmission bottleneck sizes are defined as the number of viral particles transmitted from a donor to a recipient host that successfully establish genetic lineages within the recipient. Quantifying the magnitude of these bottlenecks is important for understanding the ecological and evolutionary dynamics of viruses at multiple scales, as these bottlenecks bridge processes occurring at the between-host and within-host levels (Zwart and Elena, 2015; McCrone and Lauring, 2018). At the population-level, tight transmission bottlenecks can act to slow down the rate of viral adaptation, as beneficial mutations that arise within a donor host can be lost during transmission to a recipient host (Abel et al., 2015; Zaraket et al., 2015; Zwart and Elena, 2015; Geoghegan et al., 2016). However, they may also be advantageous to a viral population, for example by facilitating the population’s path through a rugged fitness landscape and by purging cheaters (Zwart and Elena, 2015). At the within-host level, tight transmission bottlenecks lead to lower levels of viral genetic diversity in recipient hosts and genetic drift playing an important role in shaping the viral population during the early stages of a recipient’s infection (Gutiérrez et al., 2012; Abel et al., 2015; Zwart and Elena, 2015; McCrone and Lauring, 2018; McCrone et al., 2018). Finally, quantifying transmission bottleneck sizes is important for more applied reasons: having estimates of the bottleneck size may help determine whether it is possible to reconstruct who-infected-whom in an outbreak setting and will point towards inference methods that might be the most suitable to use (Hall et al., 2016; Campbell et al., 2018; Duault et al., 2022).

Several statistical methods have recently been developed to estimate transmission bottleneck sizes from viral deep sequencing data (Zwart and Elena, 2015; Emmett et al., 2015; Sobel Leonard et al., 2017; Ghafari et al., 2020). All of these approaches rely on first characterizing the genetic variation that is present in both the donor and the recipient of an identified transmission pair. They then restrict their analyses to the subset of sites that are polymorphic in the donor. One approach (the presence/absence method) estimates bottleneck sizes by asking which of the variants identified in the donor are also detected in the recipient and which are not. This approach does not yield precise estimates of the transmission bottleneck size *N*_*b*_ and can underestimate *N*_*b*_ under certain circumstances (Sobel Leonard et al., 2017). A second approach (the binomial sampling method) instead makes use of variant frequencies quantified in the recipient, rather than just their presence or absence. However, it assumes that the observed differences in variant frequencies between a donor and a recipient arise from the process of viral sampling alone (Emmett et al., 2015; Poon et al., 2016). This approach can also lead to underestimates of the transmission bottleneck size (Sobel Leonard et al., 2017). A third approach (the betabinomial sampling method) similarly makes use of variant frequencies from the recipient but additionally accounts for deviations between donor and recipient variant frequencies that arise from demographic noise during the early period of exponential viral growth in the recipient (Sobel Leonard et al., 2017). Finally, a haplotype-based approach to transmission bottleneck size estimation has been developed (Ghafari et al., 2020); it extends the betabinomial sampling method to account for genetic linkage between loci.

Applications of these inference methods to viral sequence data have indicated that transmission bottlenecks are tight for many viral pathogens. Several studies have estimated bottleneck sizes of 1-3 viral particles for plant viruses (Moury et al., 2007; Betancourt et al., 2008; Sacristán et al., 2011). Tight transmission bottlenecks of 1-5 viral particles have also been estimated for human viruses, including influenza viruses (McCrone et al., 2018; Valesano et al., 2020), HIV-1 (Keele et al., 2008), and most recently SARS-CoV-2 (Martin and Koelle, 2021; Nicholson et al., 2021; Braun et al., 2021a,b; Wang et al., 2021; Lythgoe et al., 2021; Li et al., 2022; Bendall et al., 2023). When bottlenecks are tight, as in these cases, there is little genetic diversity that is transferred from a donor to a recipient. For acute infections, with little time to accrue new mutations, this often times leads to overall low levels of viral diversity in infected hosts. When there is no viral genetic diversity observed in a donor sample, estimation of transmission bottleneck size is not possible for that transmission pair. Studies that estimate bottleneck sizes (such as the ones cited above) therefore often rely on combining data from across a large number of transmission pairs to quantify an average bottleneck size. Within experimental settings, barcoded viruses can be used to increase host genetic diversity and thereby to improve resolution of transmission bottleneck sizes (Varble et al., 2014; Amato et al., 2022). However, natural settings do not afford us with this possibility.

There are two other issues to consider when using only variants identified in donors for transmission bottleneck size estimation. One issue is that the time of the infectious contact is not known in many cases, and the donor is unlikely to be sampled exactly at the point of transmission. If variant frequencies change rapidly over the course of infection (as observed in longitudinal studies of acute influenza and SARS-CoV-2 infections (McCrone et al., 2018; Valesano et al., 2020; Popa et al., 2020)), estimates of bottleneck size that are based on the frequencies of donor-identified variants are likely to be considerable underestimates. A second issue is that existing methods all assume that viral particles that found the infection in the recipient are randomly sampled from the donor. However, it could be the case that genetically similar virions are aggregated and transmit together, as would be the case with collective infectious units (Sanjuán, 2017). If this is the case, one would erroneously infer bottleneck sizes to be tight when they might in reality be loose.

Here, we develop an approach for estimating transmission bottleneck sizes that instead makes use of *de novo* genetic variation that is observed in a recipient. Our approach adopts several of the same assumptions as the existing betabinomial sampling approach and haplotype-based extension of this approach. Specifically, it assumes that all observed genetic variation is neutral and that the viral population in the recipient host undergoes stochastic exponential growth. It differs from existing approaches, however, in that it uses a different subset of sites for inference, namely sites that are monomorphic in both the donor and recipient but carry different alleles. Consideration of these sites, rather than sites that are polymorphic in the donor, circumvents the two issues described above. To introduce our approach, we first describe the stochastic model that we assume underlies the process of viral population expansion in a recipient. We then describe the inference framework and test our approach on simulated data, showing that it accurately recovers transmission bottleneck sizes. Finally, we apply our approach to data from influenza A virus and SARS-CoV-2 transmission pairs, confirming previous findings of tight transmission bottlenecks for these respiratory viruses.

## Methods

### The stochastic within-host model

We model the dynamics of the viral population within a recipient using a multitype branching process model. The types in this model correspond to different viral genotypes. Because we assume that all mutations are neutral, each type has the same overall offspring distribution. More specifically, we assume a geometric offspring distribution, consistent with the offspring distribution under a stochastic birth-death model. The geometric distribution is parameterized with a success probability of *p*_*geom*_, where *p*_*geom*_ = 1*/*(*R*_0_ + 1) and *R*_0_ is the within-host basic reproduction number. As such, the expected number of offspring a given viral particle leaves is given by *R*_0_. The number of mutations that occur during the production of a viral offspring is assumed to be Poisson-distributed with mean *μ*. When one or more mutations occur during the production of an offspring, the resultant offspring becomes a new type. As such, we assume infinite sites. Offspring inherit the mutations of their parent and any additional mutations that may have occurred during their own birth. Because we model the virus population as asexually reproducing, genetic linkage across the virus genome is complete. Because we are interested in characterizing transmission dynamics between infections, we consider only the supercritical case corresponding to a within-host basic reproduction number of *R*_0_ *>* 1.

The virus population starts with an initial population size of *N* viral particles, which stem from the donor’s virus population. As discussed in more detail later, *N* is related, but not equivalent, to the transmission bottleneck size *N*_*b*_. All *N* initial viral particles harbor zero *de novo* mutations. These particles could in principle be genetically distinct from one another; however, none of them carry mutations that have accrued in the recipient. Any genetic variation that is present in these particles stems from the donor. We refer to viral particles without *de novo* mutations (including these *N* initial viral particles) as wild-type particles, while remaining cognizant that these could differ from one another genetically.

Under this branching process model, we can lay out all of the possible dynamic outcomes. The first possible outcome is that the virus population in the recipient goes stochastically extinct (Figure 1A). This would result in the recipient remaining uninfected and (necessarily, but trivially) zero mutant lineages successfully establishing in the recipient. The second possible outcome is that the wild-type viral lineage, seeded by the *N* initial particles, establishes (Figure 1B). In this case, there will be an infinite number of mutant lineages that will successfully establish. This is because the wild-type viral population will ultimately grow geometrically at rate *R*_0_*e*^−*μ*^, and each of the wild-type viral particles in the ever-growing population may give rise to a mutant lineage that will also establish in the viral population. The third possible outcome is that the wild-type viral lineage goes extinct but a single mutant lineage, seeded by a wild-type viral particle, establishes (Figure 1C). Finally, the fourth possible outcome is that the wild-type viral lineage goes extinct but that two or more mutant lineages, seeded by two or more wild-type viral particles, establish (Figure 1D). In the case of a successful infection (Figures 1B-1D), the overall viral population will grow geometrically at rate *R*_0_ once the population has reached a large size.

**Figure 1.**
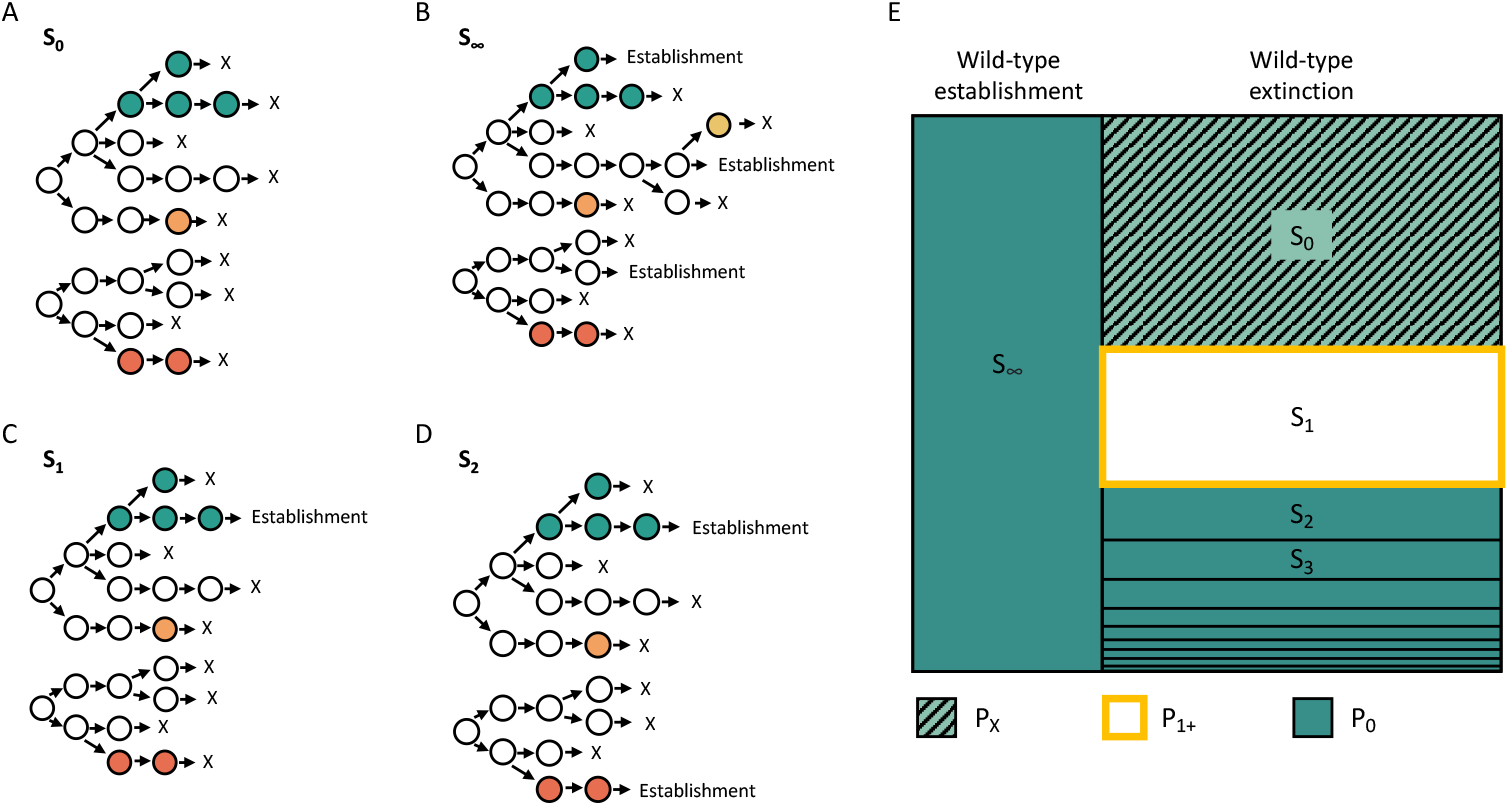
Possible dynamic outcomes in the recipient. (A) The viral population in the recipient may go stochastically extinct, leading to no infection in the recipient. (B) The wild-type viral lineage may successfully establish. (C) The wild-type viral lineage may go stochastically extinct after generating a single mutant lineage that successfully establishes. (D) The wild-type viral lineage may go stochastically extinct after generating two or more mutant lineages that successfully establish. Here, we show a case of two mutant lineages successfully establishing. Outcomes (B)-(D) result in successful infection of the recipient. Wild-type particles are shown in white. Mutant lineages are shown in different colors. In (A)-(D), *N* = 2 wild-type viral particles start off the infection in the recipient host. (E) Summary of possible dynamic outcomes. For each outcome, *S*_*l*_ denotes the number of mutant lineages *l* that establish. Outcomes are color-coded by the number of clonal variants *k* that would be observed under the outcome. The portion of the outcome space labeled *P*_*X*_ denotes the probability that the viral population in the recipient goes extinct. The portion of the outcome space labeled *P*_0_ denotes the probability that zero clonal variants establish in the recipient’s viral population. The portion of the outcome space labeled *P*_1+_ denotes the probability that at least one clonal variant establishes in the recipient’s viral population.

For a given outcome, we can quantify the number of variants that arose and fixed in the viral population of the recipient. We refer to these variants as *clonal* variants. In the case of the viral population going extinct (the first outcome; Figure 1A), the infection in the recipient did not establish and we will not have observed this outcome in a transmission pair. We refer to the probability of this outcome as *P*_*X*_ . In the case of the wild-type viral lineage establishing (the second outcome; Figure 1B), the number of clonal variants will be zero, because none of the mutations that arose in any of the mutant lineages will fix. In the case of the wild-type viral lineage going extinct but successfully seeding two or more mutant lineages prior to extinction (the fourth outcome; Figure 1D), the number of clonal variants will similarly be zero, because none of the mutations that arose in any of the mutant lineages will fix under an infinite sites assumption. Finally, in the case of the wild-type viral lineage going extinct but successfully seeding a single mutant lineage prior to extinction (the third outcome; Figure 1C), the number of clonal variants will be at least one. It will be exactly one if only a single mutation occurred during the generation of the mutant lineage and no additional clonal variants arose in this mutant lineage. It will be greater than one if more than one mutation occurred during the generation of the mutant lineage and/or if additional mutations occurred in this mutant lineage that ultimately fixed. Figure 1E graphically summarizes all of these possible dynamic outcomes.

### Derivation of the probability distribution for the number of clonal variants

The multi-type branching process model, resulting in the different possible outcomes shown in Figure 1, contains 3 parameters: the initial wild-type viral population size *N*, the within-host basic reproduction number *R*_0_, and the per genome, per infection cycle mutation rate *μ*. Here, we are specifically interested in estimating the initial viral population size *N* . Estimates of *N* will be used to calculate the transmission bottleneck size *N*_*b*_, as discussed in more detail below. To estimate *N*, we need to ask, for a given recipient harboring *k* clonal variants, what is the likelihood that the initial viral population size was *N* = 1, 2, 3, *etc*.? These likelihoods can be calculated if we can calculate the probability distribution for a recipient harboring *k* = 0, 1, 2, *etc*. clonal variants, for given values of *N, R*_0_, and *μ*. In the Supplemental Material, we derive the expression for this probability distribution, based on the different possible dynamic outcomes shown in Figure 1E.

We can confirm the accuracy of our analytical results in two ways. First, previous work by Bozic et al. (2016), in the context of cancer dynamics, derived an equation for the number of clonal variants one would expect in a population undergoing birth-death dynamics, given an initial population size of *N* = 1. This expected number is given by *δu/*(1− *δ*), where their parameter *δ* corresponds to 1*/R*_0_ and their mutation parameter *u* corresponds to our *μ*, under the assumption that *μ* is small (*<<* 1). Figure 2A shows the expected number of clonal variants across a range of within-host *R*_0_ and across a range of mutation rates *μ*, as calculated from their equation. In Figure 2B, we plot the expected number of clonal variants, calculated from the probabilities of *k* = 0, 1, 2, *etc*. clonal variants we derived in the Supplemental Material, under the assumption of *N* = 1. The quantitative similarity of the plots shown in Figures 2A and 2B demonstrate the accuracy of our clonal variant derivation. In the Supplemental Material, we further show how we can derive their equation using our expressions, under the assumption of a low mutation rate.

**Figure 2.**
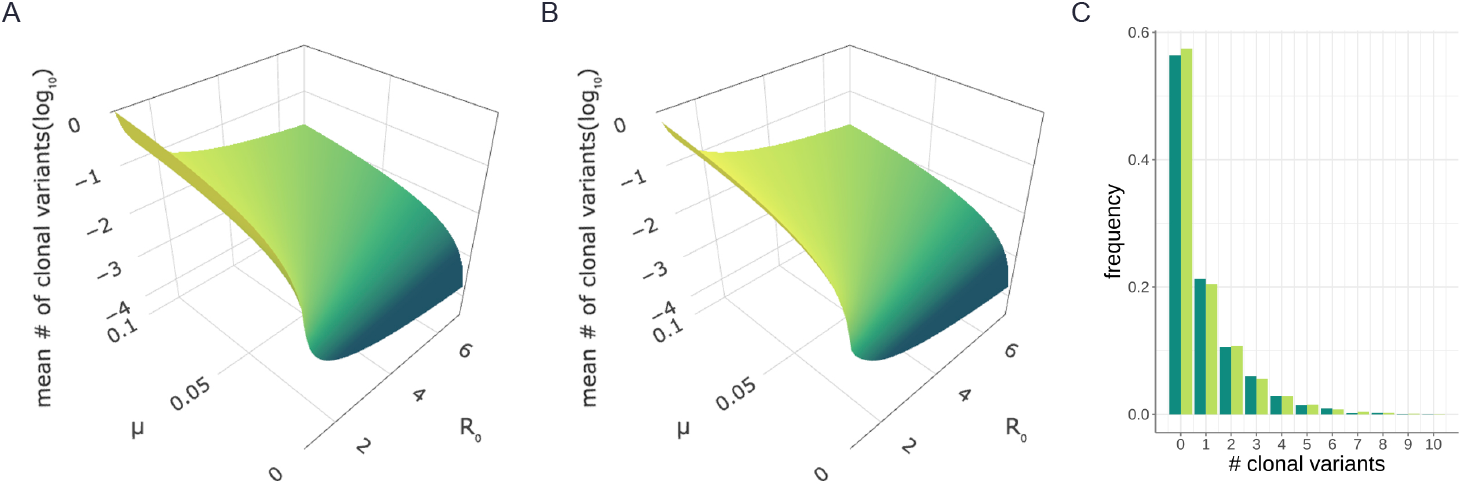
Confirmation of our analytical results. (A) The number of clonal mutations expected when *N* = 1, as derived by Bozic and coauthors using a birth-death model (Bozic et al., 2016). Mean numbers of clonal mutations are shown across a range of *R*_0_ and *μ* parameter values. (B) The mean number of clonal mutations, as calculated from our derived clonal variant probability distribution, parameterized with *N* = 1. (C) Histogram showing the proportion of simulations that resulted in *k* = 0, 1, 2, 3, *etc*. clonal variants (dark green), alongside our derived predictions (light green). Proportions were calculated using 4,000 stochastic simulations that resulted in successful infection. Simulations and analytical results shown in panel (C) were parameterized with *N* = 2, *R*_0_ = 1.2, and *μ* = 0.2.

The second way we can check our analytical results is through extensive numerical simulation of the branching process model. For a given simulation, we can determine whether the viral population went stochastically extinct or whether infection was successful. For those simulations establishing successful infection, we can determine the number of clonal variants that evolved. To check our clonal variant derivation, we plot in Figure 2C the fraction of simulations that resulted in *k* = 0, 1, 2, 3, *etc*. clonal variants from 4,000 simulations that were each parameterized with an initial viral population size of *N* = 2, a within-host basic reproduction number of *R*_0_ = 1.2, and a per genome per infection cycle mutation rate of *μ* = 0.2. Alongside this empirical distribution, we plot the analytically-derived clonal variant probabilities under this parameterization. The quantitative similarity of these distributions demonstrates the accuracy of our analytical derivations.

## Results

### Application to simulated data

Before applying our statistical method to sequence data from empirical transmission pair studies, we first applied our approach to simulated (mock) data. To this end, we simulated the branching process forward model until we obtained 100 successful recipient infections. Forward simulations were all performed with a within-host basic reproduction number of *R*_0_ = 1.6 and a mutation rate of *μ* = 0.4. Instead of assuming that the initial viral population size *N* was the same across all recipients, we assumed that the initial number of wild-type viral particles was Poisson-distributed with mean *λ* = 2.1 (Figure 3A; dark green). (Simulations with a higher *N* had a lower chance of going stochastically extinct so higher *N* simulations were overrepresented in the mock data set, which we account for, as described in greater detail below.)

**Figure 3.**
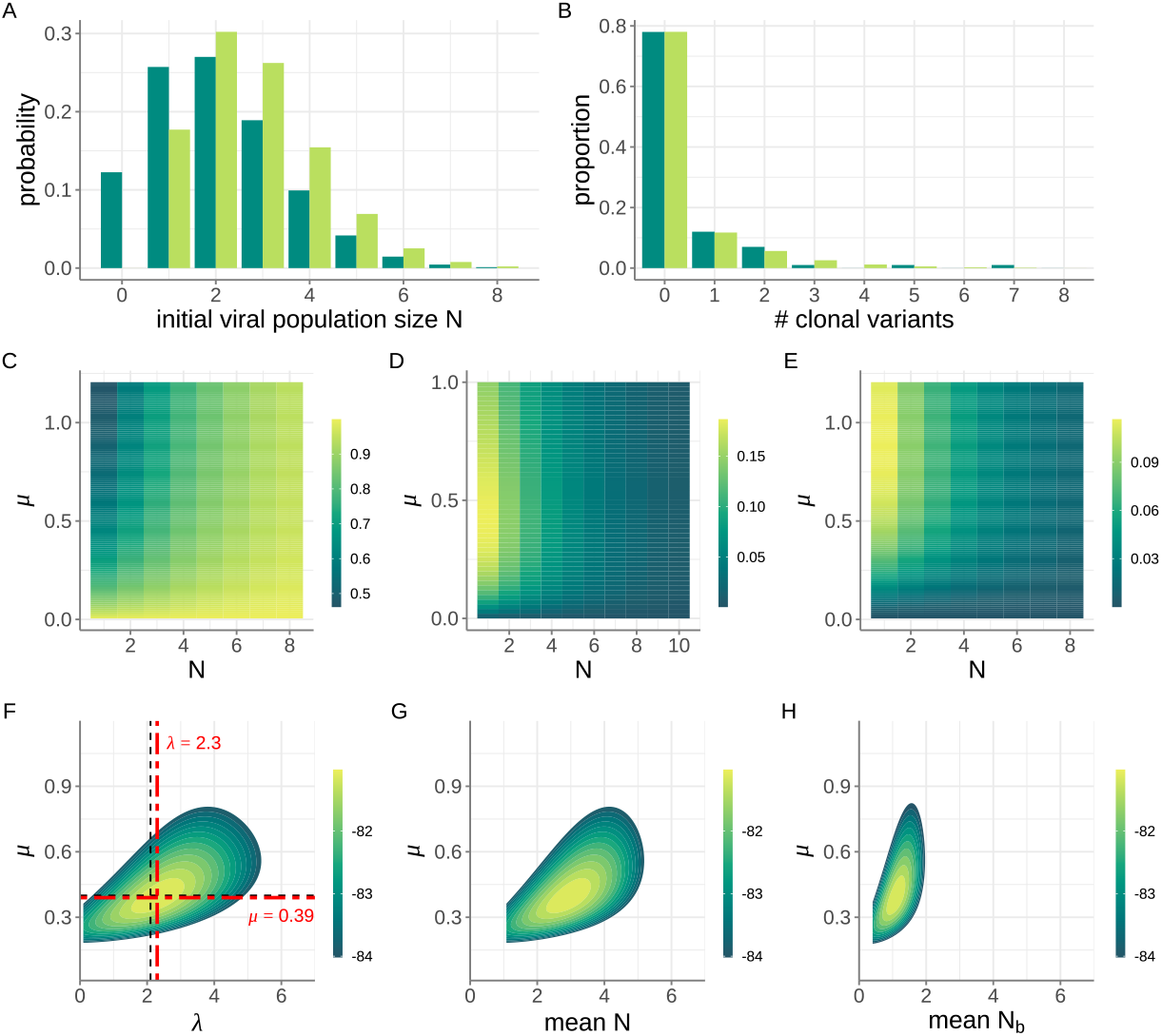
Application of our inference method to a mock dataset of 100 transmission pairs. (A) Poisson probability distribution showing the the distribution of initial viral population sizes *N* that seed potential recipient infections (dark green bars). Here, the mean of the Poisson distribution is *λ* = 2.1. The probability distribution of the initial viral population size being *N*, conditional on successful infection, is also shown (light green bars). (B) Proportion of simulated infections that resulted in *k* = 0, 1, 2, *etc*. clonal variants (dark green bars), alongside proportions predicted using the maximum likelihood values for parameters *λ* = 2.3 and *μ* = 0.39 (light green bars). Of the 100 simulated infections, 78 recipients had no clonal variants, 12 recipients had one clonal variant, 7 recipients had two clonal variants, 1 recipient had three clonal variants, 1 recipient had five clonal variants and 1 recipient had seven clonal variants. (C) Probabilities of observing *k* = 0 clonal variants across a range of *N* and *μ* values. (D) As in panel C, with probabilities of observing *k* = 1 clonal variant. (E) As in panel C, with probabilities of observing *k* = 2 clonal variants. (F) Log-likelihood plot, showing the log(probability) of observing the mock dataset given parameters *λ* and *μ*. Dashed black lines show the true values of *λ* and *μ*. Dash-dotted, bolded red lines show the maximum likelihood values of *λ* and *μ*. (G) Log-likelihood plot, as in panel F, with the results plotted as a function of 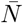 and *μ* instead of *λ* and *μ*. (H) Log-likelihood plot, as in panel F, with the results plotted as a function of 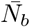 and *μ* instead of *λ* and *μ*. In (F-H), log-likelihood values are shown only for the parameter combinations that fall within the 95% confidence region.

For each of these 100 simulated successful infections, we calculated the number of clonal mutations. Figure 3B (dark green bars) shows the proportion of these 100 simulations that resulted in *k* = 0, 1, 2, 3, *etc*. clonal mutations. We then set *R*_0_ to its true value of 1.6 and attempted to jointly estimate *λ* and *μ* from this observed mock data set. To do this, we first calculated across combinations of *N* and *μ* the probability of observing *k* = 0 clonal variants (Figure 3C), *k* = 1 clonal variant (Figure 3D), *k* = 2 clonal variants (Figure 3E), *k* = 3 clonal variants (not shown), *k* = 5 clonal variants (not shown), and *k* = 7 clonal variants (not shown). (We did not perform the calculation for other values of *k* because there were no simulated infections that resulted in these other numbers of clonal variants.)

From the mock data set shown in Figure 3B (dark green bars), our goal was then to estimate *λ* and *μ* given knowledge of the within-host basic reproduction number *R*_0_. To do this, we first adjusted the Poisson distribution shown in Figure 3A (dark green bars) to reflect the distribution of initial viral population sizes we would expect across *successful* infections (Figure 3A, light green bars). This adjustment involved multiplying the Poisson probability masses by the *N* -specific probabilities of successful establishment (1 − (1*/R*_0_)^*N*^) and renormalizing. For a given transmission pair, the probability that a recipient’s viral population harbors *k* clonal variants is then given by:

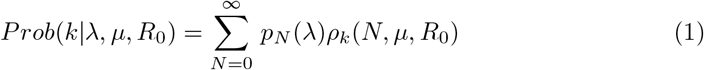

where *p*_*N*_ (*λ*) is the probability that *N* viral particles started off any given successful viral infection under an assumed Poisson distribution with mean *λ* (Figure 3A, light green bars), and where *ρ*_*k*_(*N, μ, R*_0_) is the probability that the recipient’s viral population harbors *k* clonal variants. We can calculate this probability for each of the transmission pairs in our mock data set, and then calculate the overall log-likelihood of observing the data shown in Figure 3B (dark green bars) by summing the log of these probabilities. In Figure 3F, we plot this log-likelihood surface over a broad range of *λ* values and *μ* values, while setting the within-host basic reproduction number *R*_0_ to its true value of 1.6. These results indicate that our inference approach can recover the true set of parameters (*λ, μ*) on this simulated dataset of 100 transmission pairs.

In addition to plotting out the log-likelihood landscape as a function of *λ* (and *μ*), we can plot out the same results as a function of the mean initial viral population size 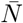 (and *μ*). The mean initial viral population size is given by:

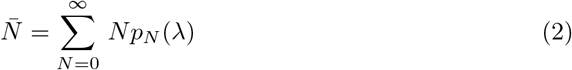

and reflects the mean of the adjusted Poisson distribution (Figure 3A, light green bars). Figure 3G plots the same log-likelihood landscape as shown in Figure 3F, with the x-axis now showing 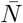 rather than *λ*. Similarly, we can plot out the same results as a function of the mean transmission bottleneck size 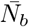 (and *μ*) (Figure 3H). The expression for the mean transmission bottleneck size is provided in the Supplemental Material.

Finally, we can use our maximum likelihood estimates of *λ* and *μ* to generate the predicted probability distribution for the number of clonal variants observed. We generate this predicted distribution using equation (1). Figure 3B shows this predicted distribution (light green bars) alongside the distribution from the simulated dataset (dark green bars). The quantitative similarity in these distribution indicates our assumption of a Poisson-distributed initial number of viral particles is consistent with patterns presented in the simulated dataset.

### Application to influenza A virus

As the first empirical application of our inference approach, we considered a rich influenza A virus (IAV) dataset from a prospective community-based cohort study (McCrone et al., 2018). The relevant portion of this dataset are 52 transmission pairs that were identified as part of this study (Supplemental Material). For each of these transmission pairs, we calculated the number of clonal variants observed in the recipient using a variant-calling threshold of 3%. (Sites with allele frequencies below 3% or above 97% were considered monomorphic.) The data consisted of 42 transmission pairs with 0 clonal variants, 5 transmission pairs with 1 clonal variant, 2 transmission pairs with 2 clonal variants, 3 transmission pairs with 3 clonal variants, and 0 transmission pairs with 4 or more clonal variants (Figure 4A). We set the within-host basic reproduction number *R*_0_ to 11.1, based on a quantitative analysis of IAV dynamics in longitudinally-studied human IAV infections (Baccam et al., 2006). We considered *λ* values between 0.01 and 4 initial viral particles and *μ* values between 0 and 3.5 mutations per genome per infection cycle.

**Figure 4.**
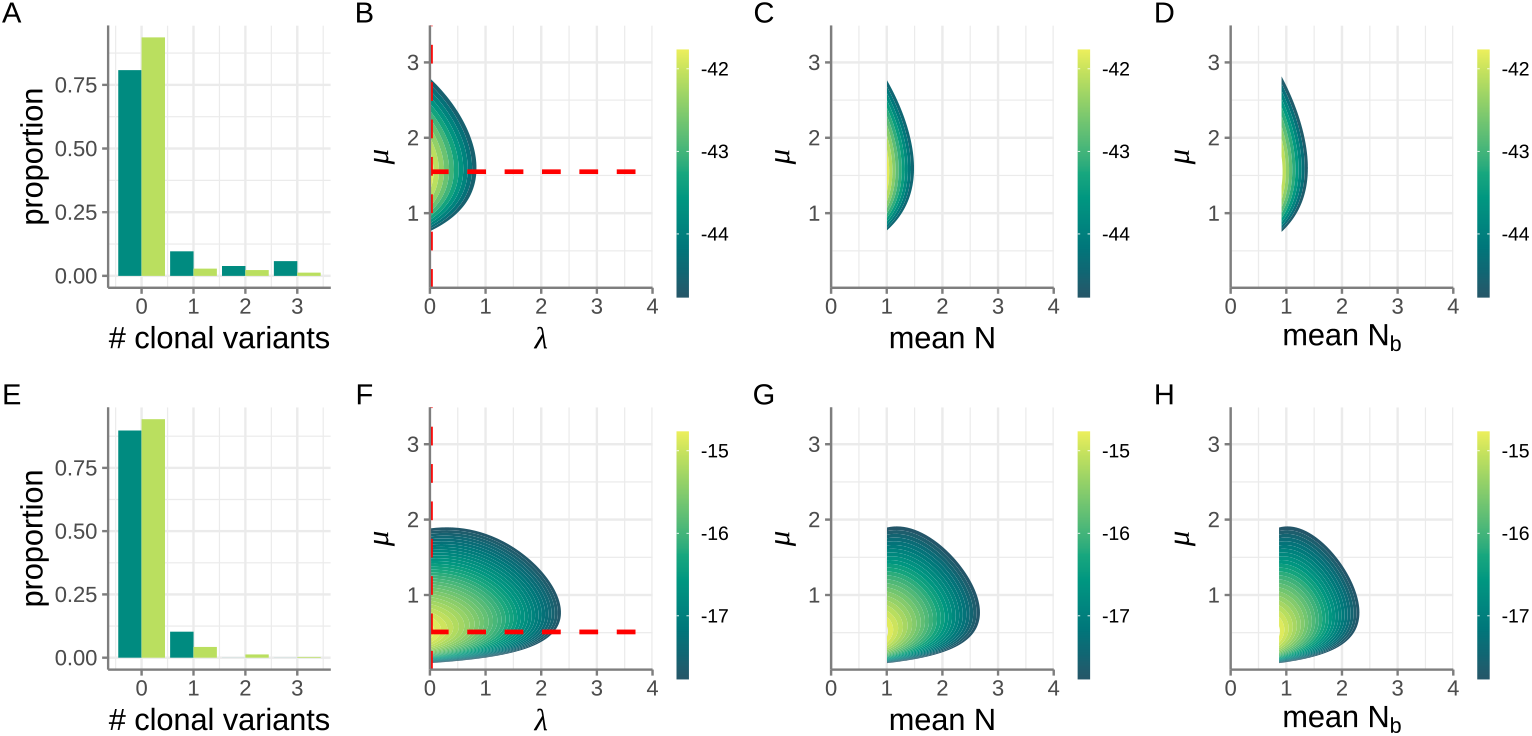
Application of our inference method to influenza A virus and SARS-CoV-2 transmission pairs. Top row shows IAV results. Bottom row shows SARS-CoV-2 results. (A) Distribution of the number of clonal variants observed across the 52 IAV transmission pairs considered. The expected distribution under the maximum likelihood estimate of *λ* = 0.01 and *μ* = 1.55 is shown alongside the empirical distribution. (B) Log-likelihood plot, showing the log(probability) of observing the IAV dataset across a range of *λ* and *μ* values. Dashed red lines show the maximum likelihood values for *λ* and *μ*. (C) Log-likelihood plot, as in panel B, with the results plotted as a function of 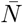 and *μ* instead of *λ* and *μ*. (D) Log-likelihood plot, as in panel B, with the results plotted as a function of 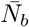 and *μ* instead of *λ* and *μ*. (E) Distribution of the number of clonal variants observed across the 39 SARS-CoV-2 transmission pairs considered. The expected distribution under the maximum likelihood estimate of *λ* = 0.01 and *μ* = 0.52 is shown alongside the empirical distribution. (F) Log-likelihood plot, showing the log(probability) of observing the SARS-CoV-2 dataset across a range of *λ* and *μ* values. (G) Log-likelihood plot, as in panel F, with the results plotted as a function of 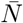 and *μ* instead of *λ* and *μ*. (H) Log-likelihood plot, as in panel F, with the results plotted as a function of 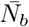 and *μ* instead of *λ* and *μ*. In panels B-D and F-H, only the log-likelihood values that fall within the 95% confidence region are shown.

Figure 4B shows the 95% confidence region for *λ* (maximum likelihood estimate = 0.01) and *μ* (maximum likelihood estimate = 1.55). Figures 4C and 4D plot these same results as a function of the mean initial viral population size 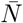 and the mean transmission bottleneck size 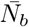, respectively. The likelihood surface shown in Figure 4D corroborates previous results of very tight transmission bottlenecks for IAV (McCrone et al., 2018). It further provides an estimate of the mutation rate that is consistent with independent mutation rate estimates based on a twelve class fluctuation test (Pauly et al., 2017). Specifically, the fluctuation test estimated the occurrence of two to three mutations on average per replicated genome. With approximately 30% of IAV mutations estimated to be lethal deleterious (Visher et al., 2016), we expect based on these results that *μ* be approximately (2 − 3) *×* 0.70 = 1.4− 2.1 mutations per replicated genome, consistent with our findings in Figures 4B-D. Finally, we used our maximum likelihood estimates of *λ* and *μ* to generate the predicted probability distribution for the number of clonal variants observed. Figure 4A shows this predicted distribution alongside the distribution from the empirical IAV dataset. The quantitative similarity in these distribution indicates that our assumption of a Poisson-distributed initial number of viral particles is consistent with patterns presented in this dataset. We do note, however, that our MLE parameter estimates appear to overestimate the proportion of clonal variants in the *k* = 0 class, and underestimate the proportion of clonal variants in higher-*k* classes.

Because our findings depend on our assumption of *R*_0_ and on the variant-calling threshold used, we re-applied our inference approach across a broader range of reasonable *R*_0_ values and across a range of different variant-calling thresholds (which can impact the number of clonal variants identified). Supplemental Figure 1 provides a sensitivity analysis of our *λ* and *μ* estimates under a range of *R*_0_ = 4.4 to 37.7 (corresponding to the minimum and maximum *R*_0_ estimates in (Baccam et al., 2006)) and under a variant-calling threshold range of 0.5% to 7%. This analysis indicates that our estimates are relatively insensitive to the exact value of *R*_0_ and the exact variant-calling threshold we use. Across the range of parameters considered, the maximum likelihood estimate of *λ* remained at 0.01 mean viral particles and the mutation rate ranged between *μ* = 0.83 mutations per infection cycle and 1.75 mutations per infection cycle. Higher variant-calling thresholds and higher within-host *R*_0_ values were associated with higher maximum likelihood estimates of *μ*. Of note, the overestimation of the probability mass in the *k* = 0 class (and the underestimation of the probability masses in the *k* ≥ 1 classes) goes away at lower *R*_0_ values, indicating that literature estimates of within-host *R*_0_ values may be high.

### Application to SARS-CoV-2

Next, we applied our inference approach to a previously published SARS-CoV-2 transmission pair dataset from Austria (Popa et al., 2020). This dataset included 39 identified transmission pairs from early on in the SARS-CoV-2 pandemic (spring 2020)(Supplemental Material). Based on shared genetic variation between donors and recipients, transmission bottlenecks sizes were estimated to be tight (Martin and Koelle, 2021; Nicholson et al., 2021), on the order of 1-3 viral particles. Here, we reanalyzed these same transmission pairs using our new inference approach, again using a variant-calling threshold of 3%. The data consisted of 35 transmission pairs with zero clonal variants, 4 transmission pairs with one clonal variant, and 0 transmission pairs with two or more clonal variants (Figure 4E). We set the within-host basic reproduction number *R*_0_ to 7.4, based on a quantitative analysis of SARS-CoV-2 dynamics in longitudinally-studied human SARS-CoV-2 infections (Ke et al., 2021). We again considered *λ* values between 0.01 and 4 initial viral particles and *μ* values between 0 and 3.5 mutations per genome per infection cycle.

Figure 4F shows the 95% confidence region for *λ* (maximum likelihood estimate = 0.01) and *μ* (maximum likelihood estimate = 0.52). Figures 4G and 4H plot these same results as a function of the mean initial viral population size 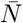 and the mean transmission bottleneck size 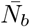, respectively. The likelihood surface shown in Figure 4H corroborates previous results of very tight transmission bottlenecks for SARS-CoV-2 (Martin and Koelle, 2021; Nicholson et al., 2021; Lythgoe et al., 2021; Braun et al., 2021b; Bendall et al., 2023). It further provides an estimate of the mutation rate that is largely consistent with an independent mutation rate estimate of 1− 5 *×* 10^−^6 per site per infection cycle (Amicone et al., 2022). This estimate translates to a per genome mutation rate of approximately 0.03-0.15 per infection cycle. Again, with approximately 30% of these mutations likely being lethal deleterious, we expect *μ* to be approximately 0.02-0.10 mutations per infection cycle. While our estimate of *μ* = 0.52 is higher than this expected range, our 95% confidence interval on *μ* extends into this range. Finally, we used our maximum likelihood estimates of *λ* and *μ* to again generate the predicted probability distribution for the number of clonal variants observed. Figure 4E shows this predicted distribution alongside the distribution from the empirical SARS-CoV-2 dataset. The quantitative similarity in these distribution indicates that our assumption of a Poisson-distributed initial number of viral particles is consistent with patterns presented in this dataset. We again note, however, that our MLE parameter estimates appear to overestimate the proportion of clonal variants in the *k* = 0 class, and underestimate the proportion of clonal variants in higher-*k* classes.

To determine the sensitivity of our findings to our assumption of *R*_0_ = 7.4 and the variant-calling threshold of 3% we used, we again re-applied our inference approach across a broader range of reasonable *R*_0_ values and across a range of different variant-calling thresholds. Supplemental Figure 2 shows our results under a range of *R*_0_ = 2.6 to 14.9 (corresponding to the minimum and maximum *R*_0_ estimates in Ke et al. (2021)) and under a variant-calling threshold range of 0.5% to 7%. Our results again indicate that our estimates are relatively insensitive to the exact value of *R*_0_ used and the exact variant-calling threshold used. As was the case with our IAV analysis, estimates of *μ* were higher at the higher variant-calling threshold and at higher within-host *R*_0_ values. Again, the overestimation of the probability mass in the *k* = 0 class (and the underestimation of the probability masses in the *k* ≥ 1 classes) is less at lower *R*_0_ values, again indicating that literature estimates of within-host *R*_0_ values may be high.

## Discussion

Here, we developed a new statistical approach for estimating transmission bottleneck sizes from viral deep sequencing data from donor-recipient transmission pairs. This approach differs from previous approaches in that it does not use the subset of viral sites that are identified as polymorphic in the donor. Instead, our approach relies on the number of clonal variants observed in the recipient. Observed clonal variants arise *de novo* shortly after transmission and are particularly well suited for estimating bottleneck sizes when bottlenecks are likely to be tight.

Our approach carries several advantages over existing approaches. First, transmission pairs where the donor does not show any genetic variation are still informative and can be included in our analysis. Second, a misspecification of donor versus recipient in a transmission pair does not impact results, as the number of clonal variants is the same with a correct donor/recipient assignment or the reverse. Third, existing studies that have looked at longitudinal viral samples have indicated that variant frequencies are highly dynamic over the course of an acute infection, consistent with a small within-host effective population size. As such, variant frequencies from a donor sample that are used to estimate bottleneck sizes may not reflect variant frequencies present in the donor at the time of transmission, and would lead to underestimates of *N*_*b*_. Even if bottleneck sizes were large, changes in variant frequencies due to genetic drift in recipients would similarly bias *N*_*b*_ estimates to be low. In contrast, our approach does not rely on variant frequencies in a donor, nor does it rely on variant frequencies in a recipient. As such, it is not subject to these same biases. Examination of our datasets also indicates that clonal variants remain clonal over the course of a recipient’s infection, such that the timing of the sampling event does not impact our dataset and thus does not impact our bottleneck size estimates. Finally, if viral particles from a donor are not randomly sampled, this does not impact our inference, while it would again bias *N*_*b*_ estimates to be low with existing inference approaches.

Despite these advantages of our new inference approach, there are some limitations to it. First, estimation of transmission bottleneck sizes requires more than a single transmission pair. Second, our approach depends on a very limited subset of the donor and recipient deep sequencing data. However, for the reasons we described above (dramatic changes in variant frequencies over the course of acute infections and the possibility of non-random sampling of the donor’s viral population), we do not believe that patterns of shared genetic variation between the donor and the recipient can be particularly informative of transmission bottleneck sizes. Our approach does ignore *de novo* genetic variation that is subclonal in the recipient, however (that is, *de novo* variants that are called in a recipient but are not fixed). Our approach could, in principle, be extended to accommodate these variants. However, based on longitudinal analyses of IAV infections (McCrone et al., 2018), we also think that many of these subclonal variants come and go over the course of an infection, such that they are not informative of the transmission bottleneck size, but instead are more informative of the extent of genetic drift that occurs over the course of an acute infection. We thus do not recommend extension of our approach to accommodate subclonal variants. Additional limitations of our approach include our assumption of infinite sites and our assumption that all genetic variation is neutral. We do not believe that the infinite sites assumption would substantially bias our results because the number of clonal variants is very small compared to the length of the viral genome. We also do not believe that the neutrality assumption would substantially bias our results in the case of small transmission bottleneck sizes because genetic drift dominates in this regime, such that small fitness differences between viral particles will not impact the viral population’s evolutionary dynamics. Lethal deleterious mutations will simply act to lower the mutation rate estimate or decrease the effective *R*_0_ of the viral population.

In our application to influenza A virus and to SARS-CoV-2, we found that transmission bottleneck sizes were very tight, consistent with previous findings of small *N*_*b*_. This raises the question of what environmental and molecular mechanisms constrain transmission bottleneck sizes. Are the number of viral particles that reach the respiratory tract of a recipient limited? Or do many viral particles reach a recipient’s respiratory tract but host and/or viral factors limit the number of viral lineages that establish? Our results of tight transmission bottleneck sizes for IAV and SARS-CoV-2 also indicate that reductions in viral population sizes between transmission events will have a large impact on shaping these viruses’ patterns of evolution and adaptation at the population level. Will these small bottlenecks ultimately act to impede viral adaptation or to facilitate it? And how will these tight bottlenecks impact population-level viral patterns, including patterns of antigenic change, genetic diversification, and deleterious mutation loads? Addressing these questions through theoretical and empirical studies will facilitate our understanding of viral transmission dynamics and ultimately guide our ability to curb the spread of these infectious diseases.

## Supplemental Information

### Derivation of the probability distribution for the number of clonal variants

Here, we derive the probabilities associated with the possible dynamic outcomes shown in Figure 1E and then show how these probabilities can then be used to obtain the probability distribution for the number of clonal variants. Consider first a contact between a donor and a recipient that results in the transfer of *N* infectious viral particles. Following this transfer, the within-host dynamics of the viral population depend on the within-host basic reproduction number *R*_0_.

Because the initial viral population size may be small, we consider the viral population dynamics to be subject to demographic stochasticity. More specifically, we assume that the viral population undergoes stochastic birth-death dynamics. This is dynamically equivalent to a branching process model with a geometric offspring distribution parameterized with a success probability of *p*_*geom*_ = 1*/*(*R*_0_ + 1), where *R*_0_ is the mean number of offspring produced (Lloyd-Smith et al., 2005). When *R*_0_ *<* 1, this corresponds to the subcritical case, and the viral population will die out with probability 1. In terms of the possible dynamic outcomes in Figure 1E, this corresponds to *P*_*X*_ = 1, regardless of the initial viral population size *N* . When *R*_0_ *>* 1, this corresponds to the supercritical case. In this case, the viral population still has the possibility of going stochastically extinct. This occurs with probability:

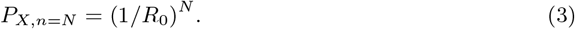

The probability of the viral population dying out *P*_*X*_ is the same as the probability of there being no mutant lineages that establish in the viral population (*S*_0_), and corresponds to the scenario depicted in Figure 1A.

Because *P*_*X*_ = 1 in the subcritical case of *R*_0_ *<* 1, we consider the remaining dynamic outcomes shown in Figure 1E only in the supercritical case. We can next derive the probability that the wild-type viral lineage successfully establishes (labeled *S*_*∞*_), corresponding to the scenario depicted in Figure 1B. To calculate this probability, we have to consider the rate at which mutations occur during replication in the viral population. We assume a per genome, per infection cycle mutation rate *μ*, with the number of mutations occurring during the production of a viral progeny being Poisson-distributed with this mean. Given this assumption, the probability that a mutation does not occur during the production of a viral progeny is given by *e*^−*μ*^ and the probability that one or more mutations occur during the production of a viral progeny is given by 1 -*e*^−*μ*^. Given this set-up, we can decompose the overall geometric offspring distribution into two separate offspring distributions: that of offspring that are genetically identical to the parent and that of offspring that differ from the parent by at least one mutation (Supplemental Figure 3A). We define the wild-type offspring distribution as the distribution of offspring from wild-type individuals that are themselves wild-type. We further define the mutant offspring distribution as the distribution of offspring from wild-type individuals that instead carry *de novo* mutations. The wild-type offspring distribution is given by a negative binomial distribution with parameters *r*_*w*_ = *e*^−*μ*^ and success probability *p*_*geom*_. The mutant offspring distribution is given by a negative binomial distribution with parameters *r*_*m*_ = 1 − *e*^−*μ*^ and success probability *p*_*geom*_. If the number of wild-type offspring from a wild-type particle is *X ∼ NB*(*r*_*w*_, *p*_*geom*_) and the number of mutant offspring from a wild-type particle is *Y ∼ NB*(*r*_*m*_, *p*_*geom*_), the overall offspring distribution from a wild-type particle is *X* + *Y∼ NB*(*r*_*w*_ + *r*_*m*_, *p*_*geom*_) = *NB*(1, *p*_*geom*_) = *Geom*(*p*_*geom*_), thereby recovering the assumed overall offspring distribution.

Now that we have the wild-type offspring distribution defined, we can calculate the probability that the wild-type viral lineage establishes. This is given by:

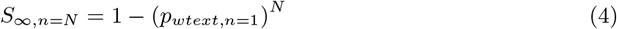

where *p*_*wtext,n*=1_ is the probability that a wild-type viral lineage, starting with a single infectious viral particle (*n* = 1) goes extinct. If the mean number of wild-type offspring (given by *R*_0_*e*^−*μ*^) exceeds one, this probability can be calculated numerically using equation (4) in Nishiura et al. (2012), using the parameters of the wild-type offspring distribution (*r*_*w*_ and *p*_*geom*_). If the mean number of wild-type offspring is less than one, *p*_*wtext,n*=1_ = 1, and *S*_*∞,n*=*N*_ = 0.

Now that we have derived *S*_*∞,n*=*N*_ and *S*_0,*n*=*N*_, we turn to deriving *S*_1,*n*=*N*_, *S*_2,*n*=*N*_, *S*_3,*n*=*N*_, etc. Deriving these probabilities is a bit more involved. Our general approach is to first calculate the final size distribution of wild-type particles, conditional on wild-type lineage extinction, and then to use this final size distribution to generate a probability mass function for the number of mutant lineages generated from the wild-type viral population. From this distribution, we calculate the probability mass function for the number of mutant lineages that establish. This probability mass function gives us *S*_0,*n*=*N*_, *S*_1,*n*=*N*_, *S*_2,*n*=*N*_, etc., thus completing our analytical expressions for the possible dynamic outcomes shown in Figure 1E.

To first calculate the final size distribution of wild-type particles, we use previous analytical results derived in Nishiura et al. (2012) and Blumberg and Lloyd-Smith (2013b). For both the wild-type supercritical case (*R*_0_*e*^−*μ*^ *>* 1) and the wild-type subcritical case (*R*_0_*e*^−*μ*^ *<* 1), the final size distribution of the wild-type viral population, starting with one single infectious particle, is given by equation (1) in Blumberg and Lloyd-Smith (2013a) for a branching process model with a negative binomial offspring distribution (here, parameterized with *r*_*w*_ and *p*_*geom*_). For the subcritical case, the probability masses of this distribution add up to 1. For the supercritical case, the probability masses of this distribution add up to *p*_*wtext,n*=1_. We can thus normalize this latter distribution such that its probability masses add up to 1 (thereby conditioning on wild-type viral lineage extinction). We define this normalized probability distribution as *f*_*n*=1_. Making use of *f*_*n*=1_, we can then calculate the final size distribution of wild-type particles starting with *N* viral particles (*f*_*n*=*N*_) through recursion:

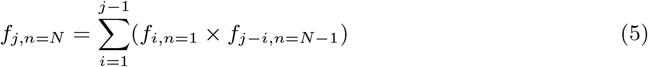

with the terminal case being *f*_*j,n*=1_. Here, *f*_*j,n*=*N*_ refers to the probability mass of the final size of wild-type particles being *j*, given an initial viral population size of *N* . Supplemental Figure 3B shows the probability mass function *f*_*n*=*N*_ .

Once we have the wild-type final size distribution *f*_*n*=*N*_ (conditional on wild-type extinction), we can calculate the probability mass function for the number of mutant lineages that were *generated* directly from the wild-type population prior to its extinction. We can do this calculation using the mutant offspring distribution. Because the mutant offspring distribution is a negative binomial distribution with parameters *r*_*m*_ and *p*_*geom*_, if the final size of the wild-type population was *j*, then the number of mutant offspring from this wild-type population itself follows a negative binomial distribution with parameters (*j × r*_*m*_) and *p*_*geom*_. We can thus iterate over all possible wild-type final population sizes to calculate an overall distribution for the number of mutant viral lineages that are generated from this wild-type population that ultimately goes extinct. We define this overall distribution as *g*_*n*=*N*_ (Supplemental Figure 3C). Finally, we can use *g*_*n*=*N*_ to calculate the distribution of the number of mutant lineages that *establish*, which we refer to as *h*_*n*=*N*_ . Each mutant lineage has a probability of establishing of 1 − 1*/R*_0_, such that:

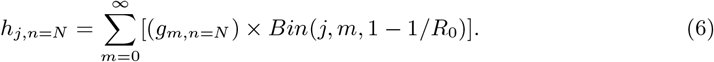

for *j* = 0, 1, 2, *etc*. (Supplemental Figure 3D). Here, *Bin*(*j, m*, 1 − 1*/R*_0_) is the binomial probability of observing *j* successes, given *m* trials and a probability of success of 1 − 1*/R*_0_. We can then calculate *S*_0,*n*=*N*_, *S*_1,*n*=*N*_, *S*_2,*n*=*N*_, etc., from *h*_*n*=*N*_ by:

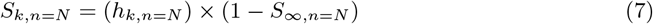

where *k* = 0, 1, 2, *etc*. (but not *∞*). This completes our analytical derivations for the outcomes depicted in Figure 1E.

As shown in Figure 1E, the probability of an infection going stochastically extinct in a recipient is given by:

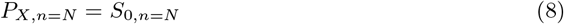

The probability of there being zero clonal variants observed in a recipient is given by:

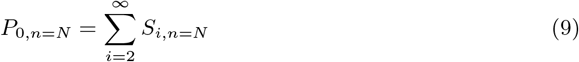

The probability of there being 1 or more clonal variants is given by:

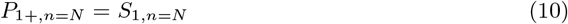

These three probabilities are shown in Supplemental Figure 3E. We now turn to resolving *P*_1+,*n*=*N*_ into the probability of observing exactly *k* = 1, 2, 3, *etc*. clonal variants.

Given the mutation rate *μ*, the probability that exactly *k* mutations occurred during the production of the mutant lineage is given by:

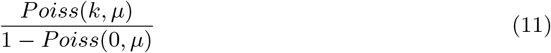

where *Poiss*(*k, μ*) is the Poisson probability of observing *k* mutations given a mutation rate of *μ*. To a first approximation, we can therefore write:

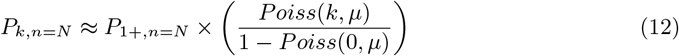

for *k* ≥ 1. This expression captures the possibility that more than one clonal variant arises during the generation of a mutant lineage from a wild-type particle. However, this expression is an approximation because there is a possibility that additional clonal variants arise following the generation of this mutant lineage (that is, during its establishment). We can correct for this possibility by modifying the above expression by probabilities of these additional clonal variants arising. Specifically, we can write the probability of there being *exactly* one clonal variant as the product of there being one mutation that occurs during the generation of the first mutant lineage and there being no additional clonal variants arising in this lineage that starts off with one mutant viral particle:

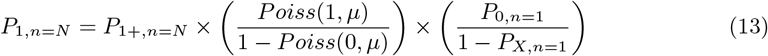

We can similarly calculate the probability of there being *exactly* two clonal variants as the probability that there are exactly two mutations that arise during the generation of the first mutant lineage and no clonal variants that arise thereafter, plus the probability that exactly one mutation arises during the generation of the first mutant lineage and exactly one clonal variant arising thereafter:

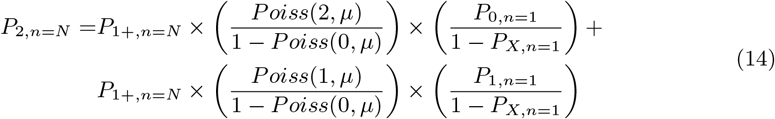

Luckily, we can directly calculate all of these terms, including *P*_1,*n*=1_, which is given by:

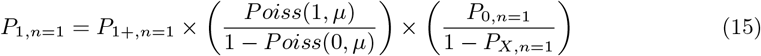

Next, the probability of observing *exactly k* = 3 clonal variants is given by the sum of the probability of 3 clonal variants arising during the generation of the first mutant lineage (and none thereafter), the probability of 2 clonal variants arising during the generation of the first mutant lineage (and 1 therefore), and the probability of 1 clonal variant arising during the generation of the first mutant lineage (and 2 thereafter). More generally, therefore, the probability of observing *exactly k* clonal variants is given by:

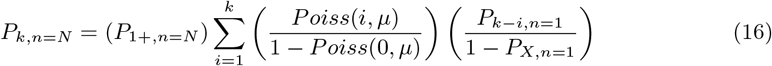

At this point, we now have *P*_*X,n*=*N*_ and *P*_*k,n*=*N*_, for *k* ≥ 0. Because we would not observe infections in recipients in the case of *P*_*X,n*=*N*_, the final probability mass distribution for the number of clonal variants observed in a recipient is given by:

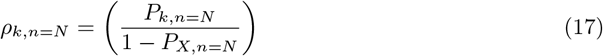

for *k* ≥ 0 (Supplemental Figure 3F).

Code for this inference approach (in both R and Matlab) is available from our GitHub site: https://github.com/koellelab/nbclonal. This site also includes an R package to calculate clonal variant probabilities.

### Rederivation of the Bozic et al. (2016) equation for the mean number of clonal variants

Bozic et al. (2016) derived an equation for the expected number of clonal variants under a scenario of a population size starting with a single individual (*N* = 1). In their work, the underlying model was a birth-death model with parameter *δ* defined as the ratio of death rate to birth rate (*d/b*) and parameter *u* defined as the probability of a mutation occurring during the production of an offspring. The equation they derived for the mean number of clonal variants is given by their equation (46): *δu/*(1 − *δ*).

Our underlying model is a branching process model, parameterized with a geometric offspring distribution. This distribution corresponds to the offspring distribution realized in a birth-death model. Our branching process model has parameter *R*_0_ defined as the ratio of birth rate to death rate (such that 1*/R*_0_ is the ratio of death rate to birth rate, *δ*). Our model also has parameter *μ*, defined as the mean number of mutations that occur at birth, with the distribution of mutations occurring at birth being Poisson distributed. The probability that zero mutations occur during the production of an offspring is thus given as *e*^−*μ*^. When *μ* is small, the probability that a (single) mutation occurs during the production of an offspring is 1 − *e*^−*μ*^, which is approximately *μ*. As such, when *μ* is small, it is approximately equal to Bozic et al.’s *u* parameter. Finally, to correspond with the assumption in (Bozic et al., 2016) of the population starting with a single individual, we set the initial viral population size *N* to 1.

With a small mutation rate *μ*, the possible outcomes shown in Figure 1E consist primarily of *S*_*∞*_, *S*_0_, and *S*_1_, and the probability that more than one clonal variant establishes within the *S*_1_ outcome is negligibly small. As such, the mean number of clonal variants is given by *S*_1_ conditional on infection, such that 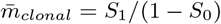. This is equivalent to 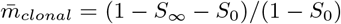 As detailed above, *S*_0_ = 1*/R*_0_ when *N* = 1. When *μ* is small, we can approximate *S*_*∞*_ as 1 − 1*/*(*R*_0_*e*^−*μ*^) = 1 − 1*/*(*R*_0_(1 − *μ*)). Substituting, the numerator becomes: (1 − *S*_*∞*_ − *S*_0_) = *μ/R*_0_ = *δu*, and the denominator becomes (1 − *S*_0_) = 1 − 1*/R*_0_ = 1 − *δ*. As such, the expected number of clonal variants from our analytical expressions, parameterized with *N* = 1, agree with equation (46) provided in Bozic et al. (2016).

### Calculation of the mean transmission bottleneck size 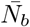

Then mean bottleneck size is given by:

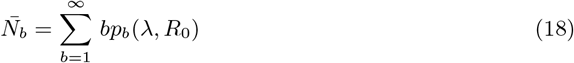

where *p*_*b*_(*λ, R*_0_) denotes the probability that the bottleneck size is *b* in a successful infection. In turn, *p*_*b*_(*λ, R*_0_) is given by:

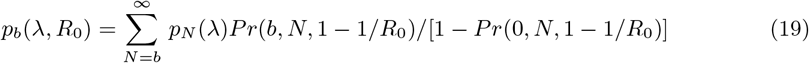

where *Pr*(*b, N*, 1 − 1*/R*_0_) is given by the binomial probability that exactly *b* out of the *N* initial viral particles successfully leave genetic lineages in the recipient host. This binomial probability is calculated using a success probability of 1 − 1*/R*_0_ for each initial viral particle.

### Quantification of the number of clonal variants for the influenza A virus dataset

We applied our inference approach to a previously published IAV transmission pair dataset from Michigan, USA (McCrone et al., 2018). The raw sequencing data have previously been made available by the authors (SRA BioProject: PRJNA412631). Individuals in the same household were considered transmission pairs if they displayed symptoms within 7 days of one another, were both positive for the same IAV subtype, and were infected with viruses that were genetically more similar to one another that 95% of epidemiologically unlinked pairs based on the L1-norm. Within these pairs, donors and recipients were established based on which individual displayed symptom onset first with at least a day of symptom onset between them. Based on these criteria, McCrone et al. (2018) narrowed 124 putative household transmission events down to 52 pairs that had sufficiently high quality sequencing data from both donor and recipient. Supplemental text file 1 provides a list of these transmission pairs and the number of clonal variants observed in each recipient, across a range of different variant-calling thresholds ranging from 0.25% to 7%. We used the datafile no cut trans freq.csv from the authors’ GitHub to calculate the number of clonal variants in this supplemental text file.

### Quantification of the number of clonal variants for the SARS-CoV-2 dataset

We applied our inference approach to a previously published SARS-CoV-2 transmission pair dataset from Austria (Popa et al., 2020). The raw sequencing data have previously been made available by the authors (SRA BioProject: PRJEB39849). To establish transmission pairs, the authors combined information on intrafamily cases with information on epidemiological transmission chains. Additional telephone investigations were used to validate inferred transmission pairs. The data set consisted of 39 transmission pairs as reported in Data File S4 from (Popa et al., 2020). Sequencing reads were downloaded from the SRA and variants were called relative to Wuhan/Hu-1 (NC 045512.2) as described in (Martin and Koelle, 2021). The number of clonal variants in each recipient relative to their donor was calculated across a range of variant-calling thresholds, from 0.25% to 7% (Supplemental text file 2). (Note that we have previously analyzed this dataset using a 6% variant-calling threshold (Martin and Koelle, 2021). The reason we used this high threshold previously was to remove spurious low frequency variants that were shared across many of the samples and therefore inflated transmission bottleneck size estimates. Here, we use the variant-calling threshold to identify the sites that are monomorphic in both the donor and the recipient and therefore might be sites that harbor clonal variants in the recipient. As such, a lower variant-calling threshold yields a more conservative estimate on the number of clonal variants observed. In contrast, while a higher variant-calling threshold yields a more conservative estimate of shared genetic variation.)

Acknowledgments/Funding

This study was supported by National Institute of Allergy and Infectious Diseases (NIAID) Centers of Excellence for Influenza Research and Response (CEIRR) contract no. 75N93021C00017. Further support for this study was provided by DARPA INTERCEPT W911NF-17-2-0034 (KK) and NIH NIAID F31 AI154738 (MAM).

## Supplemental Figures and Tables

**Figure S1.**
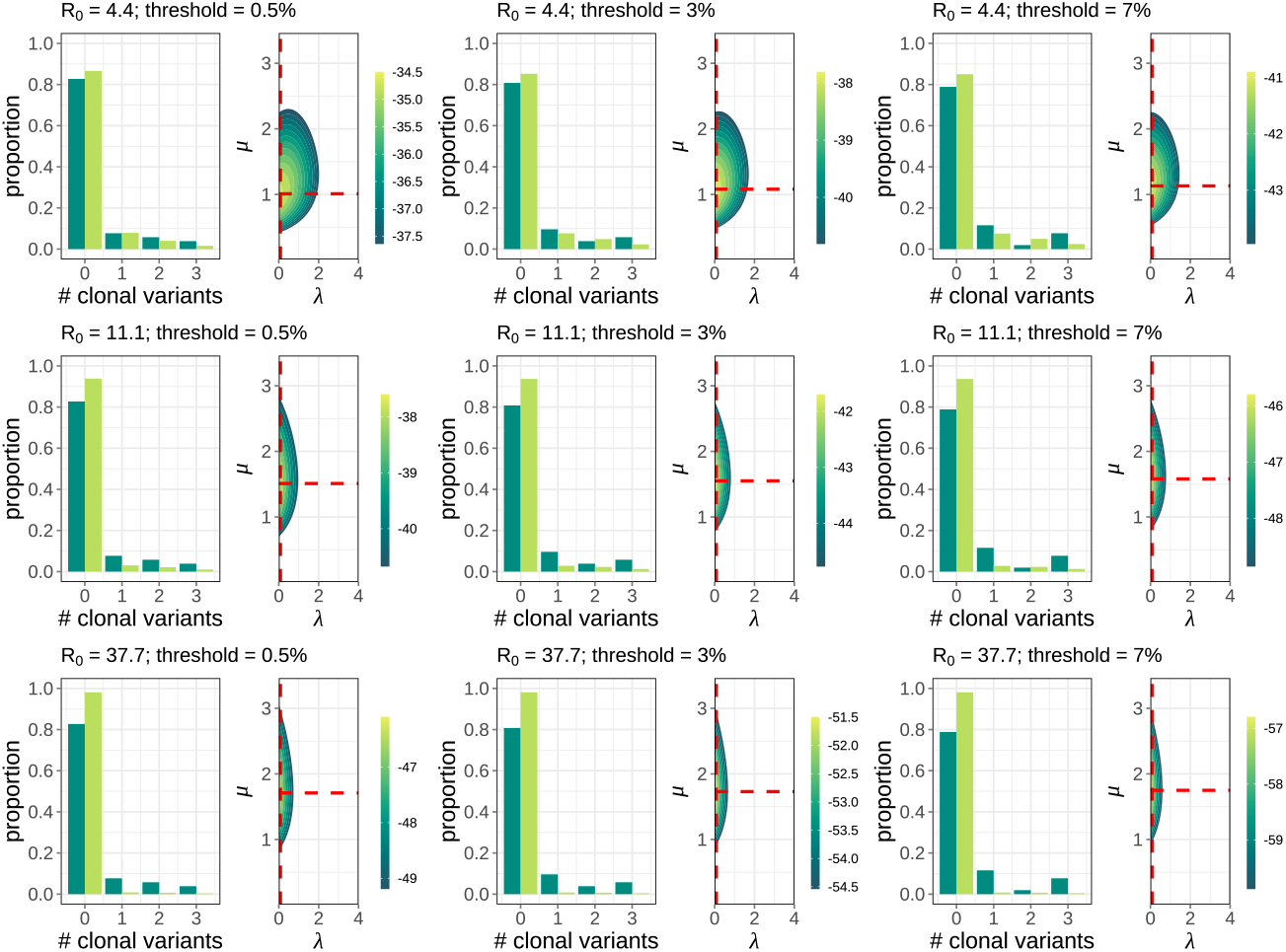
Sensitivity analysis of IAV results across different assumptions of within-host *R*_0_ and across different variant-calling thresholds. Results are shown in three rows, with the first row showing results with an assumed *R*_0_ value of 4.4, the second row showing results with an assumed *R*_0_ value of 11.1, and the third row showing results with an assumed *R*_0_ value of 37.7. Columns correspond to three different variant-calling thresholds: 0.5%, 3%, and 7%, respectively. For each *R*_0_ and threshold combination considered, we plot two panels: one similar to panel A in Figure 4, showing the empirical data and the predicted distribution using the maximum likelihood estimates and one similar to panel B in Figure 4, showing the log-likelihood plot for parameters *λ* and *μ*.

**Figure S2.**
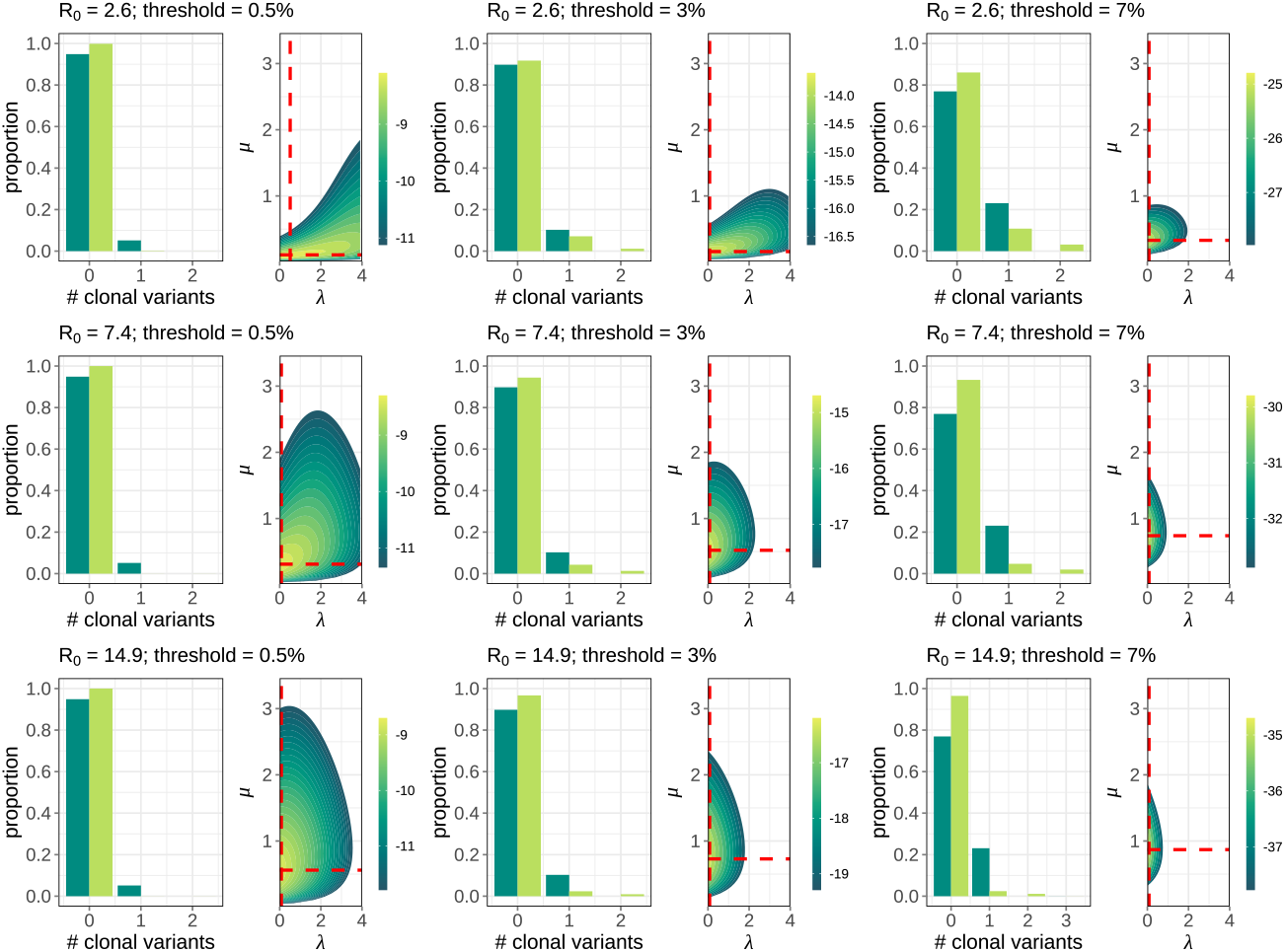
Sensitivity analysis of SARS-CoV-2 results across different assumptions of within-host *R*_0_ and across different variant-calling thresholds. Results are shown in three rows, with the first row showing results with an assumed *R*_0_ value of 2.6, the second row showing results with an assumed *R*_0_ value of 7.4, and the third row showing results with an assumed *R*_0_ value of 14.9. Columns correspond to three different variant-calling thresholds: 0.5%, 3%, and 7%, respectively. For each *R*_0_ and threshold combination considered, we plot two panels: one similar to panel E in Figure 4, showing the empirical data and the predicted distribution using the maximum likelihood estimates and one similar to panel F in Figure 4, showing the log-likelihood plot for parameters *λ* and *μ*.

**Figure S3.**
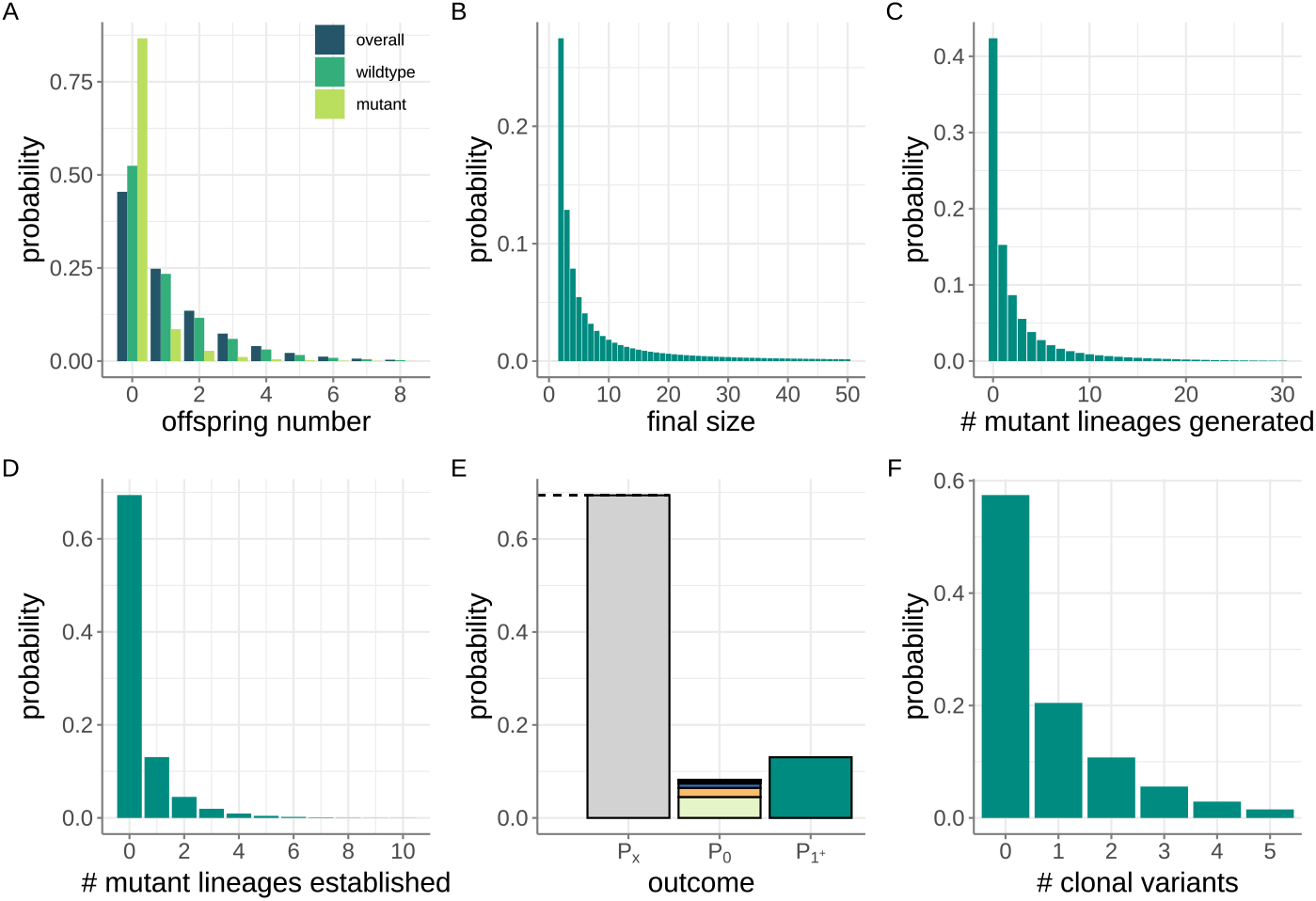
Steps involved in the derivation of the probability mass function for the number of clonal variants. (A) The overall offspring distribution, shown alongside the wild-type offspring distribution and the mutant offspring distribution. Here, the overall offspring distribution is a geometric distribution with mean *R*_0_ = 1.2 (such that *p*_*geom*_ =0.4545). The wild-type offspring distribution and the mutant offspring distribution are both negative binomial distribution with *r*_*w*_ and *r*_*m*_ calculated using a mutation rate of *μ* = 0.2. (B) The final size distribution of wild-type particles, conditional on wild-type lineage extinction. (C) The probability mass function for the number of mutant lineages generated by the wild-type viral population, conditional on wild-type lineage extinction. (D) The probability mass function for the number of mutant lineages that successfully establish, conditional on wild-type lineage extinction. (E) Calculated probabilities of the overall viral population going extinct (*P*_*X*_, as calculated by *S*_0_), of the viral population establishing with zero clonal variants (*P*_0_), and of the viral population establishing with one or more clonal variants (*P*_1+_). Dashed black line shows the analytical calculation of *P*_*X*_ via the expression given by equation (3), indicating agreement with the calculated value of *S*_0_. (F) Probability mass function for the number of clonal variants that establish in a recipient who becomes successfully infected.

**Table S1**. List of donor-recipient transmission pairs from McCrone et al. (2018) used in the bottleneck size analysis shown in Figure 4.For each transmission pair, the number of clonal variants observed in the recipient is given across a range of variant-calling thresholds (0.5%-7%). Variant-calling thresholds of 0.5%, 3%, and 7% were used in the sensitivity analysis shown in Figure S1.

**Table S2**. List of donor-recipient transmission pairs from Popa et al. (2020) used in the bottleneck size analysis shown in Figure 4. For each transmission pair, the number of clonal variants observed in the recipient is given across a range of variant-calling thresholds (0.5%-7%). Variant-calling thresholds of 0.5%, 3%, and 7% were used in the sensitivity analysis shown in Figure S2.

## Notes

### Competing Interest Statement

The authors have declared no competing interest.

